# Gene-corrected Parkinson’s disease neurons show the A30P alpha-synuclein point mutation leads to reduced neuronal branching and function

**DOI:** 10.1101/2020.11.05.369389

**Authors:** Peter A. Barbuti, Bruno FR. Santos, Paul M. Antony, Francois Massart, Gérald Cruciani, Claire M. Dording, Lukas Pavelka, Yong-Jun Kwon, Rejko Krüger

**Author notes:** Correspondence (P.B); (R.K.). (R.K.).

## Abstract

Parkinson’s disease is characterised by the degeneration of A9 dopaminergic neurons and the pathological accumulation of alpha-synuclein. In a patient-derived stem cell model, we have generated dopaminergic neurons from an individual harbouring the p.A30P SNCA mutation and compared those neurons against gene-corrected isogenic control cell lines. We have used confocal microscopy to assess the neuronal network, specifically segmenting dopaminergic neurons and have identified image-based phenotypes showing axonal impairment and reduced neurite branching. We show using multi-electrode array (MEA) technology that the neurons carrying the endogenous p.A30P alpha-synuclein mutation are functionally impaired and identified mitochondrial dysfunction as a pathogenic cellular phenotype. We report that against gene-corrected isogenic control cell lines the neurons carrying the p.A30P SNCA mutation have a deficit and are susceptible to the mitochondrial toxin and environmental pesticide Rotenone. Our data supports the use of isogenic cell lines in identifying image-based pathological phenotypes that can serve as an entry point for future disease modifying compound screenings and drug discovery strategies.

## Introduction

Parkinson’s disease (PD) is a neurodegenerative disease with no current causative treatment. It is one of the world’s fastest growing neurological disorders^1^, with a global burden expected to reach 17 million by 2040^1,2^. PD has two defining neuropathological features; the first is the degeneration of the A9 dopaminergic neurons of the substantia nigra pars compacta (SNc). It is still unclear why specifically these neurons degenerate, clinically manifesting as a movement disorder that was first defined in 1817 as the shaking palsy by James Parkinson^3^. Reviewed elsewhere, the unique cellular architecture of these dopaminergic nigral neurons including their large complex arborisation, high bioenergetic demands and resultant oxidative stress, combined with neuroanatomical telencephalization have been hypothesised among the reasons for this specific vulnerability^4–6^. Although to provide balance, an alternative and widely-held viewpoint is that PD is a prion-like disorder^7,8^. Yet, the origin of PD pathogenesis remains unclear. The formation of intra-cytoplasmic inclusion bodies (Lewy Bodies, LBs) in the dopaminergic neurons that remain is the second defining neuropathological feature of PD. These LBs contain a high concentration of lipids, crowded organelles and are immunopositive for alpha-synuclein^9,10^, a protein found ubiquitously in neurons that is localised to the pre-synaptic terminal^11^.

Rare and highly penetrant point mutations in the *SNCA* gene, which encodes the alpha-synuclein protein leads to the autosomal dominant form of PD at: p.A53T^12^, p.A30P^13^, p.E46K^14^, p.G51D^15^, and p.A53E^16^. Increased levels of physiological alpha-synuclein caused by multiplications of the *SNCA* locus by duplication^17,18^ or triplication^19^ also lead to familial PD in a dose-dependent way, with the triplication having more severe clinical symptoms and faster disease progression than the duplication^20^. Moreover, genome wide association studies (GWAS) have identified single nucleotide polymorphism (SNP) genetic variants in *SNCA* as a risk factor in sporadic PD due to modulation of alpha-synuclein expression^21–23^.

Induced pluripotent stem (iPS) cells, first described by Yamanaka^24^ and reviewed elsewhere^25^, allows differentiated cells such as dermal fibroblasts donated from an individual harbouring a pathogenic mutation to be reprogrammed to a stem cell state, enabling the generation of patient-derived iPS cell lines that can be used for disease modelling. The widespread use of targeted genome editing tools, such as clustered regularly interspaced short palindromic repeats (CRISPR)-Cas-associated nucleases (CRISPR-Cas9)^26–28^, have enabled these patient-derived iPS cells, carrying pathogenic point mutations to be corrected, thereby allowing the exact effect of the pathogenic mutation to be assessed against its isogenic control. Comparison between edited and founder isogenic cell lines therefore corrects for the effect of inter-individual variation, which, in addition to the unknown variable of the pathogenic mutation and the limits within each cellular model^29–31^, can cloud the outcome and interpretation of the observed phenotype when assessing the mutation-only effect. Consequently, the use of isogenic cell models give much needed clarity in assessing the molecular mechanisms that underpin disease pathogenicity.

Gene editing of an iPS cell line is a technically challenging procedure and refers to editing one or more cells in a colony composed of multiple cells. Not all cells will be edited and fluorescent reporters, antibiotic resistance or a combination of both, are used to enrich and/or sort out the edited cells to generate the isogenic cell line^32–34^. Previously, we showed that with the aid of high-content screening (HCS) technology we were able to generate multiple single-cell gene-corrected patient-derived iPS cell clones from a PD patient harbouring the pathogenic p.A30P mutation in *SNCA*^35^. The advantage of having single-cell clones is that it provides the guarantee that every cell in that colony and expanded cell line is edited and gene-corrected. Without this approach, it remains quite possible that the cellular composition of an iPS colony will contain a mixture of edited and unedited cells. The risk here is that cells within a colony have different proliferation rates, therefore if the gene-edited modulates cell-cycle, cell-death or developmental mutation, the cellular composition of the colony can change over the course of the culture and repeated passaging, leading to unwanted variation in the obtained findings.

In our prior study, we generated and characterised multiple gene-corrected single-cell iPSC clonal lines and found that in comparison to the isogenic founder line, the patient-derived neurons carrying the p.A30P alpha-synuclein mutation had significantly higher *SNCA* expression than the gene corrected controls^36^. In this follow-up study, we have performed functional characterisation of two gene-corrected clones and the founder cell line, where only significant differences of both gene-corrected clones against the patient cell line will be interpreted as the pathological effect of the p.A30P mutation. In this study, we have assessed complex 3D neuronal networks, segmenting the TH neurons. We have identified that irrespective of TH number, the patient-derived neurons carrying the A30P mutation show axonal impairment with reduced neuronal branching. We assessed the neuronal connectivity using multi-electrode array (MEA) technology and find reduced neuronal function in the patient neurons carrying the p.A30P mutation in *SNCA*. Furthermore, we also find mitochondrial dysfunction, with impaired respiration at basal levels. In addition, we treated these neurons using Rotenone as an environmental toxin model for PD and found that the neurons carrying the A30P mutation in alpha-synuclein have a specific reduction in neuronal viability compared to both gene-corrected iPS-derived neurons. Together, our results support the use of isogenic cell lines to model PD pathophysiology, identifying an image-based neuronal phenotype and associated functional phenotypes from the endogenous expression of the pathogenic p.A30P *SNCA* mutation, having implications for future disease modifying screening strategies.

## Materials and Methods

### Clinical patient information

The pedigree of this family, previously published shows the affected patient (IV, 5 – black arrow)^13,37^. These affected familial individuals carry an autosomal dominant heterozygous mutation in c.88G>C *SNCA* that translates the pathogenic A30P form of the alpha-synuclein protein. The patient concerned in this study is a right-hand dominant individual with an age of disease onset at 55 years with initial symptoms of extrapyramidal rigidity and bradykinesia dominant on the right side. The time between first symptoms to diagnosis was 3 months. The patient showed/presented with a good response to L-Dopa and was treated with Deep Brain Stimulation (DBS) in 2005 due to emerging motor fluctuations. The disease duration from diagnosis to clinical examination detailed in Table 1 was 13 years with the age of assessment at 68 years. The patient had 8 years of formal education up to secondary school. The dopaminergic medication at the time of the examination in ON state: Madopar p.o. 125mg ½-1/4- ½- ½, Madopar p.o. Depot 125 mg 0-0-0-1, calculated LEDD (L-DOPA equivalent daily dose) 250mg/day. The medication with central effect at the time of examination: Venlafaxin p.o. 150mg 0-0-0-1. The patient was examined by a specialized neurologist using UPDRS score in Table 1 (N.B. This is the older version of the UPDRS not the MDS-UPDRS). The clinical phenotype of the motor and non-motor symptoms are shown in Table 2.

**Table 1:** Clinical Assessment of the A30P patient in September 2010, 13 years of disease duration since diagnosis.

| Clinical assessment | Score |
| --- | --- |
| Age at the clinical assessment | 68 years |
| Hoehn and Yahr scale | Stage 3 |
| Mini-Mental State Exam (MMSE) | 26 (equivalent to MoCA 22) <sup>62</sup> |
| Swab and England Activities of Daily Living Score | 60% |
| Beck Depression Inventory (BDI) Version I total score | 6 |
| UPDRS I total score | 1 |
| UPDRS II total score | 15 |
| UPDRS IV total score | 3 |
| UPDRS III total score (ON medication, ON Stimulator) | 28 |
| UPDRS III total score (ON medication, OFF Stimulator) | 57 |
| UPDRS III total score (OFF medication, ON Stimulator) | 35 |
| UPDRS III total score (OFF medication, OFF Stimulator) | 59 |
| UPDRS III Medication ON/OFF (Stimulation ON) 28/35 | amelioration of 20% |
| UPDRS III Stimulation ON/OFF (Medication ON) 28/57 | amelioration 51% |

**Table 2:** Clinical diagnosis of the A30P patient motor and non-motor symptoms. Severity indicators: − absent; (+) discrete/variable; + mild; ++ moderate; +++ marked.

| Motor Symptoms | Severity | Non-motor symptoms | Severity |
| --- | --- | --- | --- |
| **Bradykinesia** | ++ | **Subjective cognitive impairment** | - |
| **Rigidity** | ++ | **Anxiety** | - |
| **Resting tremor** | - | **Depression** | + |
| **Postural Instability** | ++ | **Hallucinations** | - |
| **L-DOPA response** | ++ | **Dysphagia** | ++ |
| **Motor fluctuations** | ++ |  |  |
| **Dyskinesia** | - |  |  |
| **Freezing of Gait** | + |  |  |

### Cell line

A biopsy of dermal fibroblasts was donated with informed consent at 67 years of age. The generation and characterisation of induced pluripotent stem cells (iPSCs) from the dermal fibroblasts has been described^38^ and has a unique identifier HIHDNDi001-B (https://hpscreg.eu/cell-line/HIHDNDi001-B). The generation of single-cell gene-corrected patient-derived iPS clones from the A30P patient has also been described^35^. Ethical approval was obtained from the National Committee for Ethics in Research, Luxembourg (Comité National d’Ethique dans la Recherche; CNER #201411/05).

### Maintenance of iPS, generation of NPCs and differentiation to vmDA neurons

The maintenance of induced pluripotent stem cells (iPSCs) has been previously described^38^. The generation of ventral midbrain dopaminergic (vmDA) neurons was performed according to the protocol elsewhere described^39^. The vmDA neurons were generated via a multipotent neuronal precursor cell (NPC) stage, the generation and characterisation of the NPCs and vmDA neurons used this study has been previously described^35^.

### Assessment of neuronal networks

After 35 days of directed neuronal differentiation, neurons were treated with pre-warmed Accutase^®^ (Sigma-Aldrich, St. Louis, MO, USA; A6964) for approximately 10 minutes and placed in an incubator (37 °C, 5% CO_2_). The neurons were dissociated to obtain a single-cell suspension before DMEM was added and the cells centrifuged (300 g; 5 mins). The neurons were plated at 200,000 cells/coverslip with the addition of 10 μM Rho-Kinase Inhibitor Y-27632 (10 μM; Abcam, Cambridge, UK; Ab120129). After 48 hours, the neurons were fixed and stained as previously described^38^ using primary antibodies for β-3-Tubulin (TUJ1; 1:200; BioLegend, San Diego, CA, USA; 801201) and TH (1:300; Millipore, Burlington, MA, USA; AB152)^35^. For the image acquisition, 10 to 12 Z-stack images per coverslip were acquired using Zeiss spinning disk confocal microscope from four independent directed neuronal differentiations. The custom image analysis algorithms for the morphological analysis of neurology morphology, neurite morphology and skeletonisation, nuclei segmentation and TH neuron segmentation have been previously described^40,41^.

### Assessment of cellular function

#### Electrical recording using multi-electrode arrays (MEAs)

Neurons were dissociated after 45 days of directed differentiation and plated at 200,000 cells/well into a minimum of 4 wells of a 48-well (16 electrode) CytoView MEA™ (Axion Biosystems, Atlanta, GA, USA; M768-tMEA-48W). The neurons were cultured on the plate for an additional 10 days and recorded at d55. The recording of the MEA plate has been previously described^41^. Briefly, the MEA plate was recorded using the Axion Integrated Studio (AxIS) software, version 2.1 (Axion Biosystems, Atlanta, GA, USA) on the Axion Maestro (Axion Biosystems, Atlanta, GA, USA). The plate was recorded for 5 minutes at 37 °C and 5% CO_2_. The 5% CO_2_ was obtained from a compressed gas bottle (Linde, Munich, Germany) and was regulated to the MEA at 0.2 bar. Rerecording of the MEA was performed using AxIS, software version 2.5, with a Butterworth (200Hz-3kHz) filter. The Spike Detector program was used to measure activity, the Adaptive Threshold Crossing method was used to detect crossings at 6x Standard Deviation. The neural metrics were analysed using the Neural Metric Tool, software version 2.6.1 (Axion Biosystems, Atlanta, GA, USA). The threshold for an active electrode was set to five spikes per minute.

### Assessment of mitochondrial function

#### Mitochondrial bioenergetics using Seahorse

The detection of oxygen consumption rate (OCR) and extracellular acidification rate (ECAR) was determined using an XFe96 Extracellular flux bioanalyzer (Agilent technologies, Santa Clara, CA, USA) using a mitochondrial stress test. Briefly, neurons were differentiated for 44 days of directed differentiation prior to be treated with Accutase^®^ for 5-20 minutes and seeded on the assay plate precoated 24 hours prior with 200 μg/mL Laminin (Sigma-Aldrich, St. Louis, MO, USA; L2020). Neurons were seeded (80,000 cells/well) in 80 μL, a minimum of 8 technical replicates were used with the perimeter wells avoided. Cells were left for an hour at room temperature to avoid edge effect before being incubated at 37 °C. The day prior to the assay the assay cartridge was hydrated with XF-calibrant solution and incubated in a non-CO_2_ incubator as per manufacturer instructions. On the day of the experiment homemade buffered assay media was prepared for the assay containing DMEM (#D5030), supplemented with Glucose (21.25 mM; 47829), Pyruvate (40 mg/L; P2256) and L-Glutamine (2 mM; G7513) (all reagents were acquired from Sigma-Aldrich, St. Louis, MO, USA), and pH calibrated to 7.4 following 37 °C non-CO_2_ incubation. The cells were washed twice with buffered assay media to give a final volume of 175 μL and incubated for 1 hour at 37 °C in a non-CO_2_ incubator. For the mitochondrial stress test assay, Oligomycin (1 μM; 75351), FCCP (250 nM; C2920) and Rotenone and Antimycin A (5 μM; 557368 and A8674) was used (all chemicals were acquired from Sigma-Aldrich, St. Louis, MO, USA). The experiment was designed and run using Wave 2.6.0 software (Agilent technologies, Santa Clara, CA, USA). Postexperiment, the plate was lysed using RIPA buffer and normalised to protein using the BCA assay with the perimeter wells used to contain the standards in triplicate. Post-normalisation, the experimental data was exported using the Seahorse XF Cell Mito Stress Test Report Generator. The Multi-File Seahorse XF Cell Mito Stress Test Report Generator was used to assess three biological replicates with statistical analysis performed in GraphPad Prism software version 8 (GraphPad Software Inc. La Jolla, CA, USA).

#### Assessment of neuronal viability

The CellTiter-Glo^®^ Luminescent Cell Viability Assay (Promega, Madison, WI, USA; G7570) was used according to the manufacturer instructions to determine cell viability based on the real-time quantification of ATP, which is directly proportional to the number of cells in the culture. Briefly, after 34 days of directed differentiation, the neurons were treated with Accutase^®^ for 10 minutes and seeded on a 384-well plate pre-coated 24 hours earlier with Geltrex™ LDEV-Free Reduced Growth Factor Basement Membrane Matrix (Thermo Fisher Scientific, Waltham, MA, USA; A1413201). Neurons were seeded at 25,000 cells/ well and were maintained in the plate for 5 days before treatment. Rotenone (Sigma-Aldrich, St. Louis, MO, USA; 557368) or DMSO was added using the Echo^®^ acoustic liquid handler (Labcyte Inc., San Jose, CA, USA) with a 16 hour incubation. The total luminescence was normalised to the confluence percentage using the SpectraMax i3x Multi-Mode Microplate Reader i3 (Molecular Devices^®^, San Jose, CA, USA) that was taken on the same day of experiment prior to the addition of Rotenone or DMSO. The luminescence was measured using the Cytation 5 (Bio-Tek Instruments Inc., Agilent technologies, Santa Clara, CA, USA) with a minimum of eight technical replicates used per condition. The experiment was repeated following four independent neuronal differentiations.

## Results

### Morphometric analysis of neuronal network reveals axonal pathology with specific reduction of neuronal branchpoints in the A30P patient

Previously, we had used CRISPR-Cas9 technology to gene-correct the c.88G>C, p.A30P *SNCA* mutation in the patient iPSCs^35^. Two single-cell patient-derived gene-corrected isogenic iPSC clones were fully characterised with vmDA neurons were generated, characterised, and quantified^35^. In order to perform detailed neuronal network analysis, Z-stacks were generated using fluorescently labelled antibodies specific for the neuronal marker, TUJ1 and the dopaminergic marker, Tyrosine Hydroxylase (TH). A representative image of a maximum intensity image of each cell line is shown (Fig. 1A).

**Figure 1:**
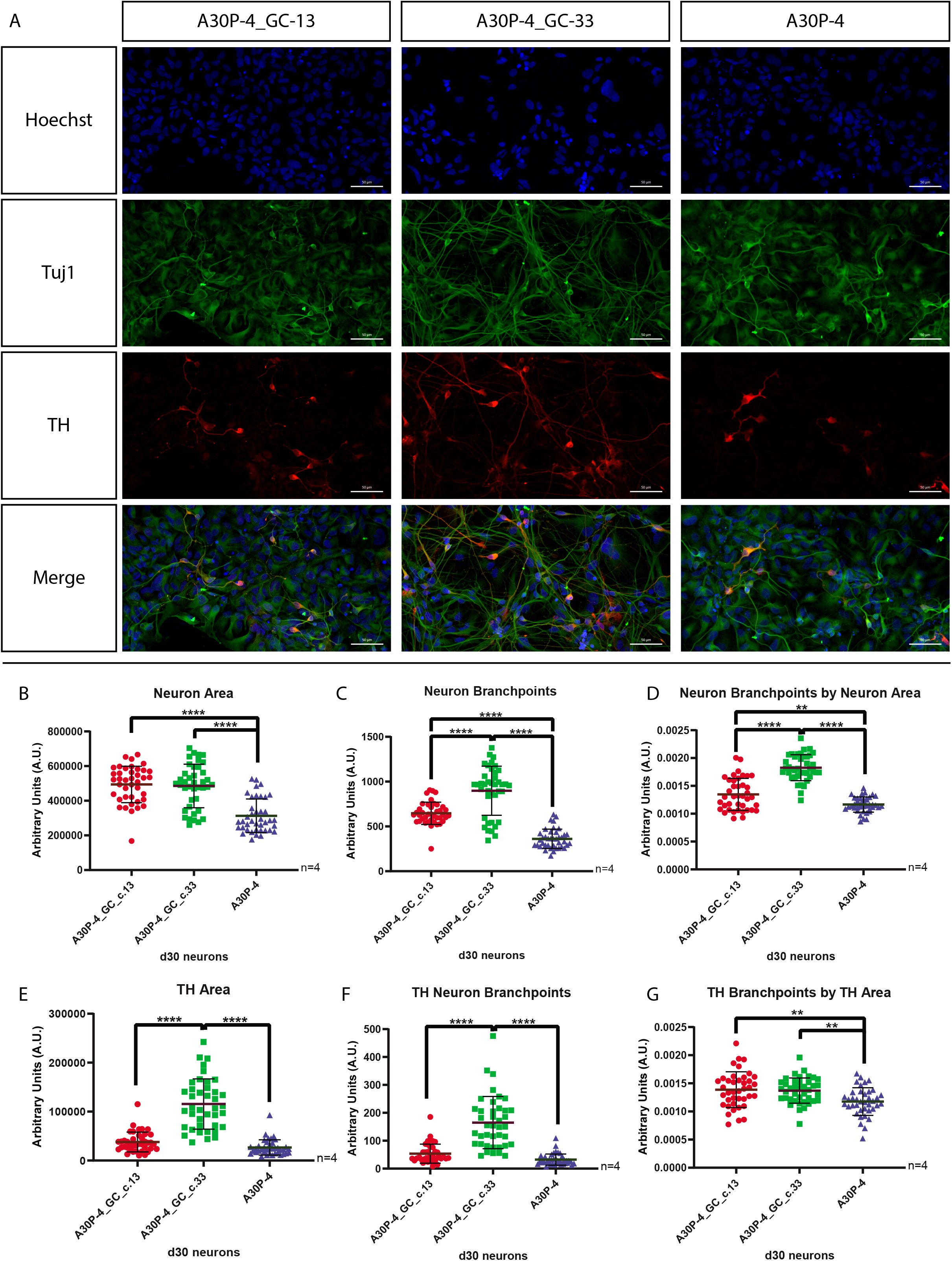
Morphometric analysis of TH network in d30 neurons. (A) Representative image of a maximum intensity projection of a Z-stack fluorescently labelled with Tuj1 and TH antibodies. Images taken using a 25x objective, scale bar is 50μM. Identification and quantification of parameters: (B) Neuron Area, (C) TH Area, (D) Neuron Branchpoints and (E) TH Neuron Branchpoints were determined. Relative normalisation resulted in (F) Neuron Branchpoints by Neuron Area, and (G) TH Branchpoints by TH Area. Four biological replicates were used for the analysis with a minimum of ten Z-stacks analysed by replicate. The graphs displayed in columns show individual values that refer to the average data per Z-stack. For all statistical analyses, an ordinary one-way ANOVA was performed with Tukey’s post-hoc multiple comparison test. All graphs were plotted as mean ± SD. ** p< 0.01, **** p< 0.0001.

Previously we found no significant difference in the number of neurons generated between the cell lines^35^. Yet, when assessing the patient-derived neurons carrying the p.A30P *SNCA* mutation against both isogenic controls our image-based analysis shows axonal pathology with neuronal area (Fig. 1B) and branchpoints (Fig. 1C) reduced. Normalisation of the neuronal branchpoints to the corresponding neuronal area confirms a reduction of neuronal branching (Fig. 1D), a newly identified pan-neuronal phenotype associated with the pathogenic p.A30P alpha-synuclein protein. In order to determine the extent of the pathology in dopaminergic neurons we performed neuronal segmentation of the TH+ neurons, representative images of neuronal and TH segmentation is shown in the supplementary material (Supp. Fig. 1.). We found significant inter-clonal variation between the two gene-corrected clones in terms of TH area (Fig. 1E) and TH neuron branchpoints (Fig. 1F) where gene-corrected clone 33 shows a significantly higher number than gene-corrected clone 13 and the founder patient line carrying the A30P mutation. However, this specific increase in TH area correlates with our previous study which found that the gene-corrected clone 33 generates approximately double the amount of TH neurons than the sister gene-corrected clone 13, and the founder patient cell line^35^. Normalising of TH branchpoints by TH area removes the clonal variation between the two gene-corrected clones due to differences in differentiation propensity (Fig. 1G). Analysis of non-TH neurons (Supp. Fig. 2) also confirms the image-based axonal phenotype of reduced neuronal branchpoints and connectivity in neurons carrying physiological levels of the pathogenic p.A30P alpha synuclein protein.

### Multi-electrode arrays (MEAs) show reduced neuronal function in A30P patient

After we had established that the neurons carrying the p.A30P alpha-synuclein mutation had a reduction in branchpoints, a measurement of synaptic connections, we hypothesised that this would influence neuronal function. We used extracellular multi-electrode arrays (MEAs) to measure neuronal activity *in-vitro*. A representative example of the seeded neurons on the MEA device is shown (Fig. 2A), where a heat-map detecting mean firing rate is the measurable parameter. An example of a raw voltage recording of a single electrode capturing a neuronal spike is shown (Fig. 2B) with a representative image of a digitally generated waveform profile displaying a neuronal action potential shown (Fig. 2C). A representative raster plot profile from a neuronal recording is displayed showing that each cell line exhibits functional active neurons (Fig. 2D). The neurons carrying the A30P mutation have reduced neuronal function compared to both gene-corrected clones with decreased mean firing rate (Fig. 2E), decreased number of synchronised spiking neurons “bursts” (Fig. 2F), and a decreased percentage of bursts (Fig. 3G). Additional parameters of neuronal function generated by the analysed neuronal recording of burst frequency, average burst duration, average number of spikes/burst, average inter-burst interval (IBI), and average IBI coefficient of variation (Figs. 2H–2L) show no difference between the neurons carrying the A30P mutation and the gene-corrected neurons without the A30P mutation.

**Figure 2:**
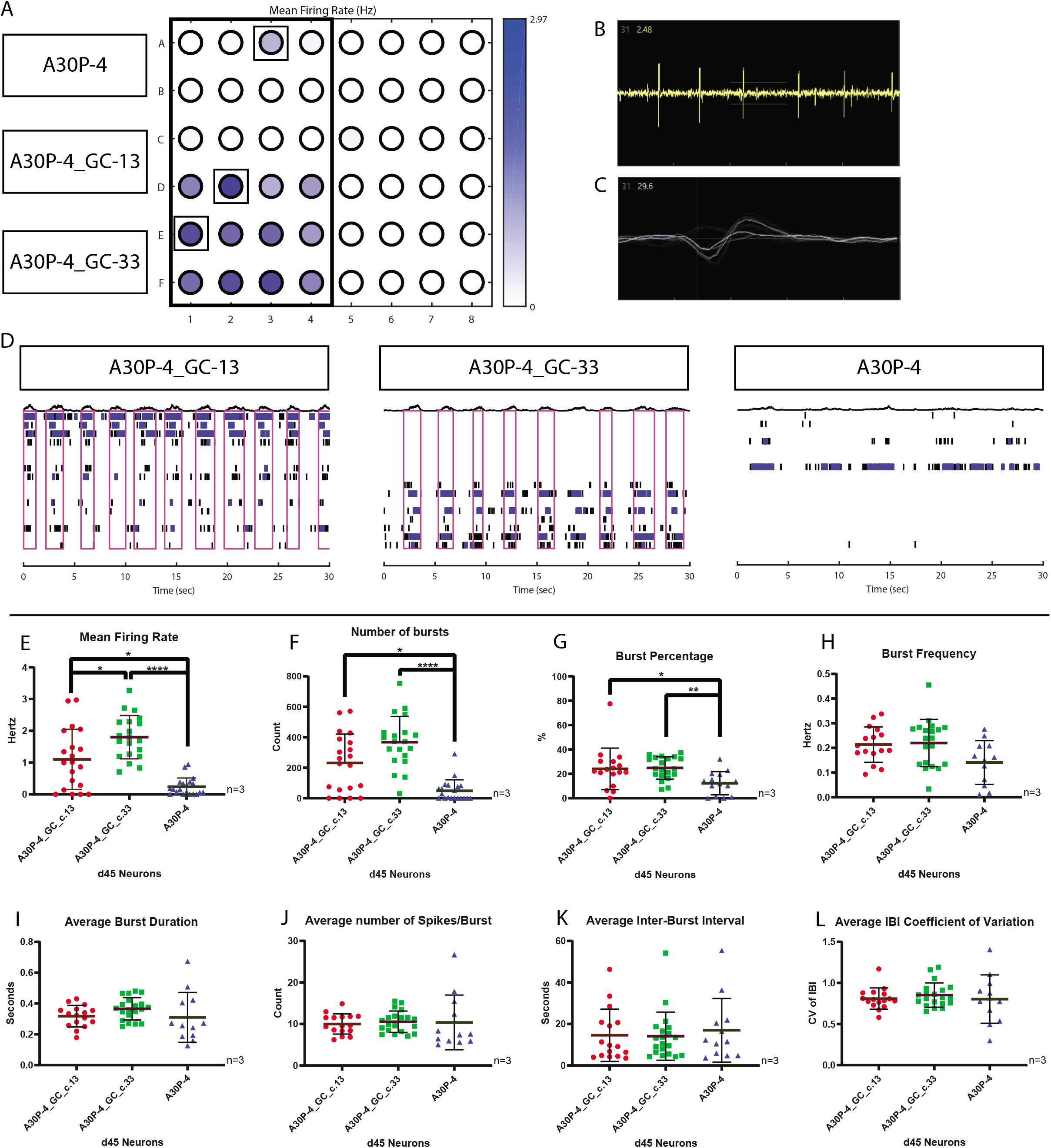
Multi-electrode array neuronal recording of d55 neurons. (A) Representative image showing the plate layout of recorded neurons with a heat map displaying the mean firing rate. (B) Representative example of a Waveform profile, (C) Representative example of a Spike plot, (D) Representative image of a raster plot profile, taken from the well highlighted in (A). Spike and burst metric parameters identified from the MEA recording: (E) Mean Firing Rate, (F) Number of Bursts, (G) Burst Percentage, (H) Burst Frequency, (I) Average Burst Duration, (J) Number of Spikes/Burst, (K) Average Inter-Burst Interval, and (L) Average Inter-Burst Interval (IBI) Co-efficient of Variation. Three biological replicates were used in the analysis with a minimum of six technical replicates per recording. The graphs displayed in columns show individual values that refer to the average data of each well, with each well composed of 16 electrodes. For all statistical analyses, a non-parametric Kruskal-Wallis test was performed using the Dunn’s post hoc multiple comparison test. All graphs were plotted as mean ± SD. *p<0.05, ** p< 0.01, **** p< 0.0001.

### Neurons carrying the p.A30P SNCA mutation have reduced mitochondrial function

The results obtained from the MEA recording show reduced neuronal function in the A30P patient neurons. Neurons are an energy intensive cell type requiring vast amounts of ATP for synaptic function that is primarily met via oxidative phosphorylation by the mitochondria. In order the determine the effect of the A30P alpha-synuclein mutation on mitochondrial respiration we used the Seahorse XFe96 extracellular flux assay to assess the cellular bioenergetics of these mutant neurons compared to the gene-corrected controls. A mitochondrial stress test was performed with direct measurement of the oxygen consumption rate (OCR). Mitochondrial stressors: Oligomycin, FCCP, Antimycin A and Rotenone were added to the neurons to assess different respiratory parameters with the merged profile plot of three biological replicates is shown (Fig. 3A). The neurons carrying the p.A30P mutation have significantly reduced mitochondrial respiration at basal levels (Fig. 3B) and reduced capability to generate ATP (Fig. 3C). There are also significant reductions in non-mitochondrial respiration (Fig. 3D) and coupling efficiency (Fig. 3E) from the p.A30P *SNCA* neurons compared to both gene-corrected isogenic control neurons.

**Figure 3:**
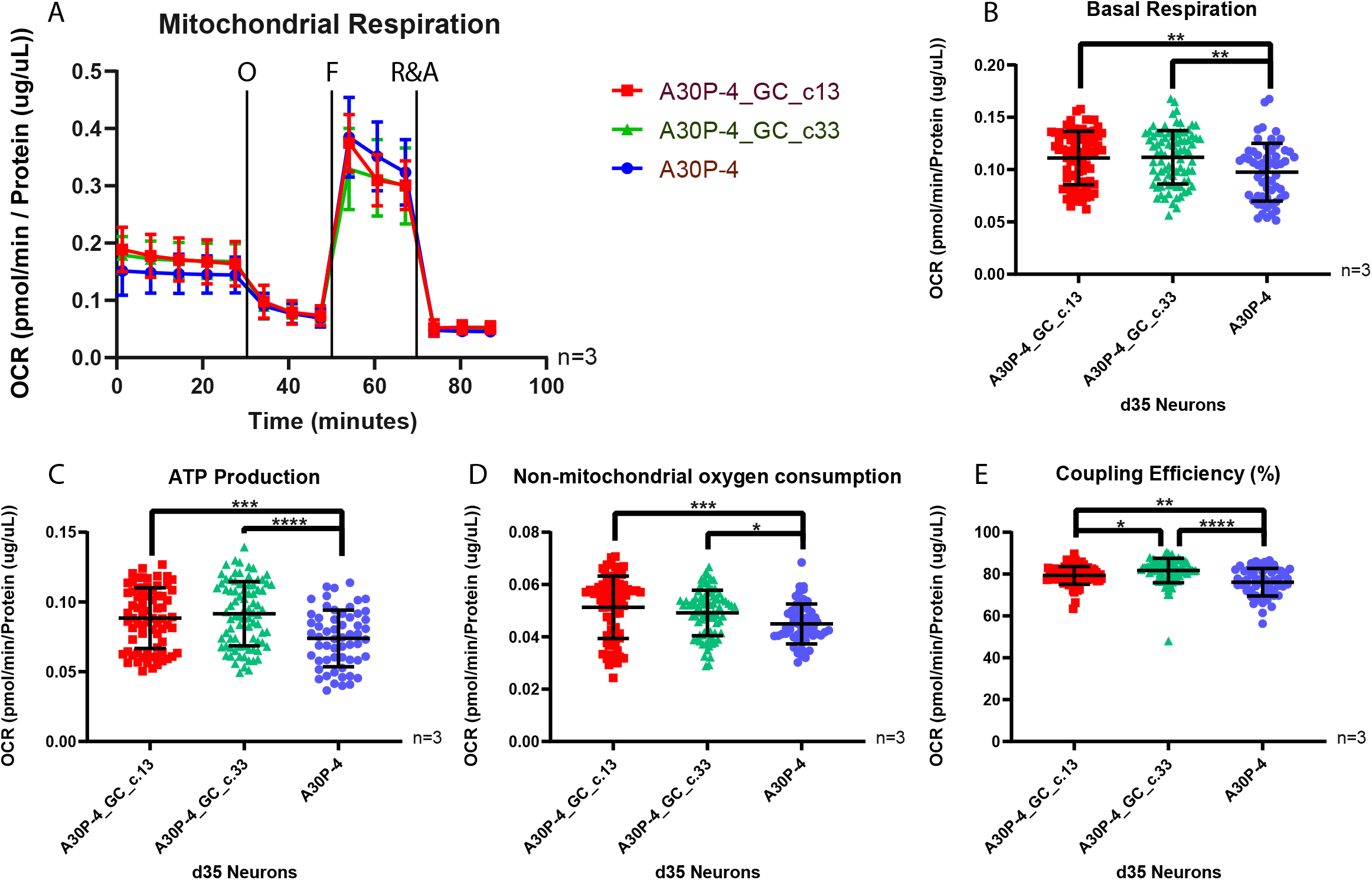
Mitochondrial dysfunction in neurons carrying a p.A30P SNCA mutation. (A) Bioenergetic profile showing oxygen consumption rate (OCR) of patient-derived neurons following a mitochondrial stress test under basal conditions and following the treatments of the ATP synthase inhibitor Oligomycin (O, 1uM), the oxidative phosphorylation uncoupler FCCP (F, 500nM), and the electron transport chain inhibitors Rotenone (Complex I) and Antimycin A (Complex III) (R&A, 10uM). The cumulative OCR profile is shown in ventral midbrain neurons differentiated for 30 days (n=3). The rates of (B) Basal Respiration, (C) ATP production, (D) Non-mitochondrial respiration and (E) Coupling efficiency are shown. For all statistical analyses, an ordinary one-way ANOVA was performed using the Tukey post-hoc multiple comparison test. All graphs were plotted as mean ± SD. *p<0.05, ** p< 0.01, ***p<0.001 **** p< 0.0001.

### Decreased neuronal viability following complex I inhibition

Having identified specific functional and mitochondrial impairments in the neurons expressing the p.A30P SNCA mutation, we hypothesised that these neurons would be specifically vulnerable to toxic insult. To assess this we used Rotenone, a complex I inhibitor that leads to the selective degeneration of TH neurons^42,43^. It is noticeable that compared to both gene-corrected clones the total luminescence, a measurement of cellular ATP in the viable neurons, although variable is significantly lower in the patient neurons carrying the A30P mutation (Fig. 4A). Treatment with increasing concentrations of Rotenone lead to a reduction in ATP and thereby a reduction in cellular viability (Fig. 4B). The neurons expressing the p.A30P SNCA mutation show a specific vulnerability to treatment with 1 nM of Rotenone leading to a significant fold decrease in cellular viability compared to both gene-corrected controls. There is no fold change in neuronal viability between patient and gene corrected controls at concentrations between 10 nM and 10 μM. At the concentration of 100 μM of Rotenone, there is a significant fold decrease in cellular viability in the neurons expressing the p.A30P SNCA mutation compared to both gene-corrected controls. Our results indicate that 100 μM of Rotenone is the concentration where the majority of neurons are no longer viable, whereas the concentration of 1 nM of Rotenone is sufficient to identify a selective fold change in cellular viability in differentially vulnerable neurons carrying a pathogenic A30P mutation in alpha-synuclein.

**Figure 4:**
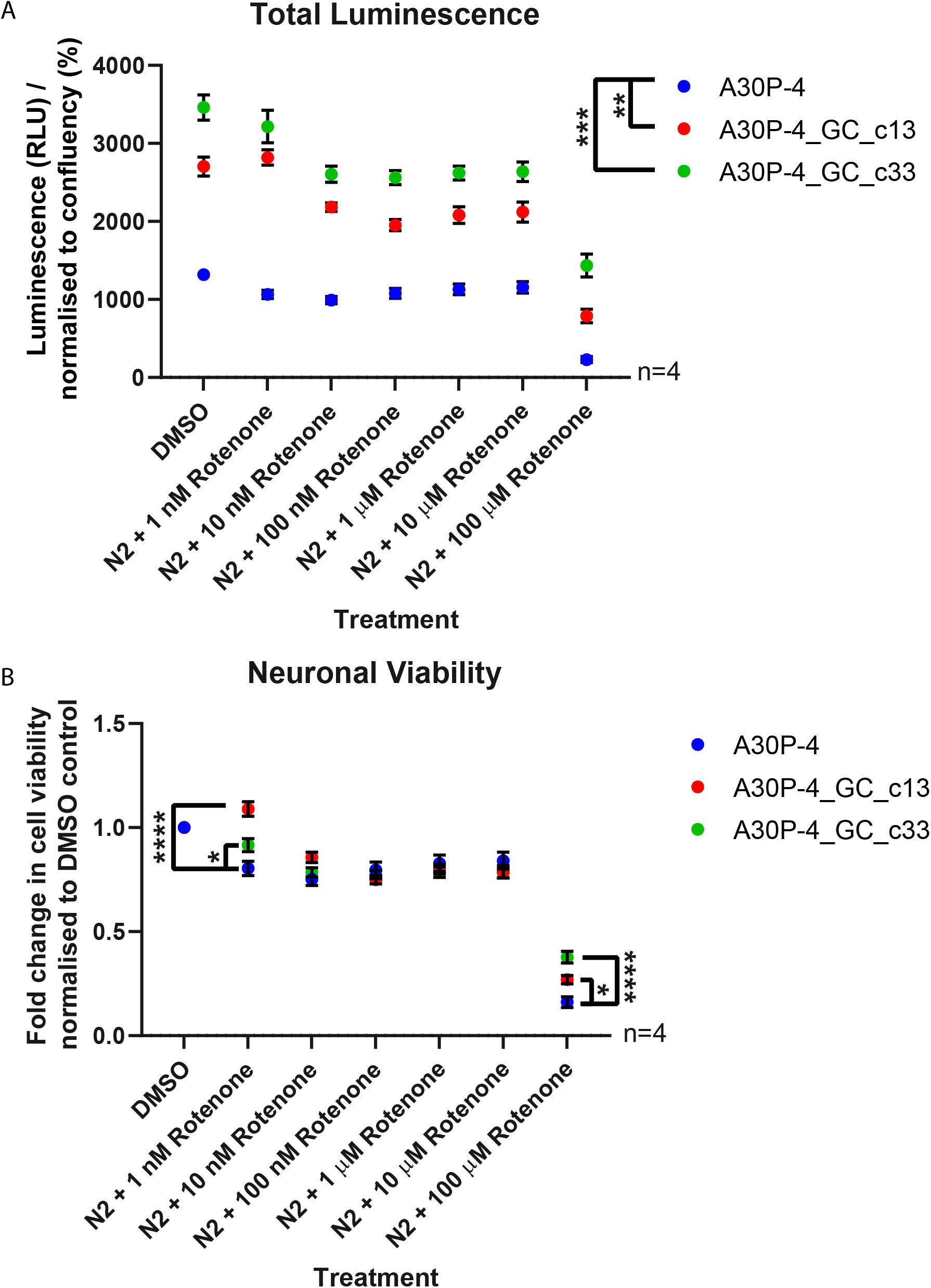
Neuronal viability after Rotenone treatment. (A) Total luminescence profile of the patient-derived neurons using the CellTiter-Glo^®^ Luminescent Cell Viability Assay with increasing concentrations of Rotenone treatment performed over 16 hours. The neurons were differentiated for 40 days, a minimum of eight technical replicates were used per analysis with four biological replicates performed. For statistical analysis, an ordinary one-way ANOVA was performed using the Tukey post-hoc multiple comparison test. (B) Fold change in cell viability of patient-derived neurons normalised to DMSO control. A two-way ANOVA was used with a Turkey’s post-hoc multiple-comparison test. All graphs were plotted as mean ± SEM. *p<0.05, ** p< 0.01, ***p<0.001 **** p< 0.0001.

## Discussion

Parkinson’s disease is a disease of neurons, with many neurons other than the innervating striatal A9 dopaminergic neurons affected^44^. However, why this specific subtype of dopaminergic neurons in the midbrain is more affected than others and why they degenerate first is a question occupying various hypotheses elsewhere reviewed^4–7,45^. Although, the physiological role(s) of alpha-synuclein is unclear and is a much debated topic^46^, one of the generally accepted physiological functions of alpha-synuclein is the regulation of synaptic vesicle endocytosis^47,48^, with synaptic dysfunction long-being associated with PD^49,50^. What we have identified from our phenotyping of functional isogenic neurons is that all neurons carrying a pathogenic point mutation in *SNCA*, expressing mutant A30P alpha-synuclein protein are impaired. Our imagebased analysis have identified that all neurons, both dopaminergic neurons and non-dopaminergic neurons show axonal impairment with reduced neuronal branching and connectivity. A general assessment of neuronal function using MEAs has also found that these pathological neurons are functionally impaired in regards to neuronal spike and burst release kinetics. Our results fit with a recent study of patient-derived isogenic dopaminergic neurons carrying the p.A53T *SNCA* mutation that found axonal pathology when compared against its isogenic control^51^. Taken together, we can consider that pathogenic alpha-synuclein facilitates axonal impairment via synaptic dysfunction. We must also consider that in the PD patient, physiological levels of pathogenic alpha-synuclein protein become symptomatic with catastrophic downstream consequences only in the sixth decade of life. Consequently, the synaptic plasticity that previously compensated for the mutation-effect must decline during the aging process and contribute to the disease pathogenesis.

Dopaminergic neurons exhibit a characteristic autonomous pacemaking activity, continuous cycles of synaptic vesicle recycling and the packaging of dopamine within those synaptic vesicles. All of which, require copious amount of energy. Improperly packaged cytosolic dopamine is subject to oxidation and can lead to increased levels of oxidant stress with pathogenic downstream affects^52^. We find that compared to both gene-corrected isogenic control cell lines, the neurons that express p.A30P alpha-synuclein have a severe energy deficit with reduced basal respiration and lower levels of ATP production per cell. Our findings align with another study that gene-corrected the patient-derived p.A53T *SNCA* mutation assessing iPS-derived dopaminergic neurons, in which deficits in mitochondrial respiration were identified as a PD phenotype^53^. Moreover, we find that the levels of cellular viability are reduced in the neurons carrying the pathogenic p.A30P alpha-synuclein variant compared to the gene-corrected isogenic controls. We have furthermore, defined and established that a concentration of 1 nM Rotenone, an environmental pesticide and well-established mitochondrial toxin specific to dopaminergic neurons^42,43^, is sufficient to significantly reduce neuronal viability in our cellular model. The identification of these neurotoxic parameters has potential implications for high-throughput disease modelling and drug discovery research by identifying screening parameters to identify vulnerable neurons, before the implementation of subsequent rescue strategies.

Our study is the first functional neuronal analysis in gene-corrected isogenic cell lines from a PD patient harbouring the pathogenic p.A30P mutation in the *SNCA* gene. Only a small number of studies have generated patient-derived gene corrected isogenic cell lines harbouring pathogenic mutations in the *SNCA* gene,^35,51,53–55^. Assessment of the endogenous p.A53T SNCA expression in dopaminergic neurons against their isogenic gene corrected control have identified mitochondrial dysfunction and apoptosis from increased basal levels of oxidative and nitrosative stress^53^, reductions in neurite length and complexity^51^, and impaired mitochondria dynamics^55^. Upregulation of ER stress and activation of the unfolded protein response pathway have been found neurons in a triplication of the *SNCA* gene dosage^54^, and we previously reported that gene-correction of the A30P mutation impairs the mitochondrial network^56^.

The importance of using isogenic patient-derived gene-corrected cell lines to model disease is that it enables assessment of the mutation-only effect in isolation from the variation within the genetic background that by itself may contribute to, or alternatively mask the mutation-specific phenotypes. However, there are still limitations within our cellular model, the typical efficiency to generate dopaminergic neurons for PD research is 10-40%^39^,^57^,^58^, although recent protocols have become more efficient^59^, these dopaminergic neurons are not “aged” and modelling age-related neurodegenerative diseases remains challenging^60^. In order to assess the effect that pathogenic mutations have on dopaminergic neurons it is fundamental to generate a higher yield of dopaminergic neurons. Successfully sorting dopaminergic neurons from a mixed neuronal culture or single-cell analysis would provide further information underlying the specific vulnerability of dopaminergic neurons. In our cellular model, we have identified that axonal impairment is not limited to TH neurons. Similarly, in a pathoanatomical study of the brain from an individual carrying the p.A30P SNCA mutation, extensive neuronal loss was not limited to the SNc, but also the locus coeruleus and the dorsal motor vagal nucleus^61^. It would be interesting to address this differential vulnerability and evaluate pure population of vulnerable neurons, i.e. dopaminergic, against a pure population of less vulnerable neurons, such as GABAergic neurons from the same individual harbouring a PD causing mutation. This would aid in identifying the specific molecular mechanisms and machinery underlying the vulnerable dopaminergic neuronal subtype that fails during PD. Furthermore, dopaminergic neurons in the SNc do not exist in isolation, improvements in cellular differentiations to regionally specific glia and microglia leading to multi-culture experiments will ultimately move patient-derived modelling closer to the patient. This point is often overlooked but is pertinent as alpha-synuclein immunopositive astroglial “coiled bodies” and oligodendroglial are found in the A30P patient brain^61^. Our findings provide the first evidence of axonal and neuronal dysfunction together with mitochondrial dysfunction as causative mechanisms that underpin PD pathology in a patient-derived isogenic neuronal model of PD harbouring the p.A30P *SNCA* mutation.

## References

1. Dorsey, E. R., Sherer, T., Okun, M. S. & Bloem, B. R. The Emerging Evidence of the Parkinson Pandemic. J. Parkinsons. Dis. 8, S3–S8 (2018).

2. Dorsey, E. R. et al. Global, regional, and national burden of Parkinson’s disease, 1990-2016: a systematic analysis for the Global Burden of Disease Study 2016. Lancet Neurol. 17, 939–953 (2018).

3. Parkinson, J. An essay on the shaking palsy. 1817. J. Neuropsychiatry Clin. Neurosci. 14, 223–36; discussion 222 (2002).

4. Bolam, J. P. & Pissadaki, E. K. Living on the edge with too many mouths to feed: Why dopamine neurons die. Mov. Disord. 27, 1478–1483 (2012).

5. Surmeier, D. J., Obeso, J. A. & Halliday, G. M. Selective neuronal vulnerability in Parkinson disease. Nat. Rev. Neurosci. 18, 101–113 (2017).

6. Diederich, N. J., Surmeier, D. J., Uchihara, T., Grillner, S. & Goetz, C. G. Parkinson’s disease: Is it a consequence of human brain evolution? Mov. Disord. 34, 453–459 (2019).

7. Brundin, P. & Melki, R. Prying into the Prion Hypothesis for Parkinson’s Disease. J. Neurosci. 37, 9808 LP – 9818 (2017).

8. Ma, J., Gao, J., Wang, J. & Xie, A. Prion-Like Mechanisms in Parkinson’s Disease. Frontiers in Neuroscience 13, 552 (2019).

9. Shahmoradian, S. H. et al. Lewy pathology in Parkinson’s disease consists of crowded organelles and lipid membranes. Nat. Neurosci. 22, 1099–1109 (2019).

10. Spillantini, M. G., Crowther, R. A., Jakes, R., Hasegawa, M. & Goedert, M. α-Synuclein in filamentous inclusions of Lewy bodies from Parkinson’s disease and dementia with Lewy bodies. Proc. Natl. Acad. Sci. 95, 6469–6473 (1998).

11. Burré, J. The Synaptic Function of α-Synuclein. J. Parkinsons. Dis. 5, 699–713 (2015).

12. Polymeropoulos, M. H. et al. Mutation in the alpha-synuclein gene identified in families with Parkinson’s disease. Science 276, 2045–7 (1997).

13. Krüger, R. et al. Ala30Pro mutation in the gene encoding alpha-synuclein in Parkinson’s disease. Nat. Genet. 18, 106–108 (1998).

14. Zarranz, J. J. et al. The new mutation, E46K, of α-synuclein causes parkinson and Lewy body dementia. Ann. Neurol. 55, 164–173 (2004).

15. Lesage, S. et al. G51D α-synuclein mutation causes a novel Parkinsonian-pyramidal syndrome. Ann. Neurol. 73, 459–471 (2013).

16. Pasanen, P. et al. Novel α-synuclein mutation A53E associated with atypical multiple system atrophy and Parkinson’s disease-type pathology. Neurobiol. Aging 35, 2180.e1–5 (2014).

17. Chartier-Harlin, M.-C. et al. Alpha-synuclein locus duplication as a cause of familial Parkinson’s disease. Lancet (London, England) 364, 1167–9 (2004).

18. Ibáñez, P. et al. Causal relation between A-synuclein locus duplication as a cause of familial Parkinson’s disease. Lancet 364, 1169–1171 (2004).

19. Singleton, A. B. et al. alpha-Synuclein locus triplication causes Parkinson’s disease. Science 302, 841 (2003).

20. Nishioka, K., Ross, O. A. & Hattori, N. SNCA Gene Multiplication: A Model Mechanism of Parkinson Disease. in (ed. Ross, O. A.) Ch. 20 (IntechOpen, 2011). doi:10.5772/24726

21. Pihlstrøm, L. et al. A comprehensive analysis of SNCA-related genetic risk in sporadic parkinson disease. Ann. Neurol. 84, 117–129 (2018).

22. Edwards, T. L. et al. Genome-Wide Association Study Confirms SNPs in SNCA and the MAPT Region as Common Risk Factors for Parkinson Disease. Ann. Hum. Genet. 74, 97–109 (2010).

23. Chiba-Falek, O., Lopez, G. J. & Nussbaum, R. L. Levels of alpha-synuclein mRNA in sporadic Parkinson disease patients. Mov. Disord. 21, 1703–1708 (2006).

24. Takahashi, K. et al. Induction of Pluripotent Stem Cells from Adult Human Fibroblasts by Defined Factors. Cell 131, 861–872 (2007).

25. Li, M. & Izpisua Belmonte, J. C. Looking to the future following 10 years of induced pluripotent stem cell technologies. Nat. Protoc. 11, 1579–1585 (2016).

26. Jinek, M. et al. A Programmable Dual-RNA-Guided DNA Endonuclease in Adaptive Bacterial Immunity. Science (80-.). 337, 816–821 (2012).

27. Mali, P. et al. RNA-Guided Human Genome Engineering via Cas9. Science (80-.). 339, 823 LP – 826 (2013).

28. Ran, F. A. et al. Genome engineering using the CRISPR-Cas9 system. Nat. Protoc. 8, 2281–2308 (2013).

29. Ferrari, E., Cardinale, A., Picconi, B. & Gardoni, F. From cell lines to pluripotent stem cells for modelling Parkinson’s Disease. J. Neurosci. Methods 340, 108741 (2020).

30. Chia, S. J., Tan, E.-K. & Chao, Y.-X. Historical Perspective: Models of Parkinson’s Disease. Int. J. Mol. Sci. 21, (2020).

31. Ke, M., Chong, C.-M. & Su, H. Using induced pluripotent stem cells for modeling Parkinson’s disease. World J. Stem Cells 11, 634–649 (2019).

32. Roberts, B. et al. Systematic gene tagging using CRISPR/Cas9 in human stem cells to illuminate cell organization. Mol. Biol. Cell 28, 2854–2874 (2017).

33. Steyer, B. et al. Scarless Genome Editing of Human Pluripotent Stem Cells via Transient Puromycin Selection. Stem cell reports 10, 642–654 (2018).

34. Arias-Fuenzalida, J. et al. FACS-Assisted CRISPR-Cas9 Genome Editing Facilitates Parkinson’s Disease Modeling. Stem Cell Reports 9, 1423–1431 (2017).

35. Barbuti, P. et al. Using High-Content Screening to Generate Single-Cell Gene-Corrected Patient-Derived iPS Clones Reveals Excess Alpha-Synuclein with Familial Parkinson’s Disease Point Mutation A30P. Cells 9, (2020).

36. Barbuti, P. et al. Using High-Content Screening Technology as a Tool to Generate Single-Cell Patient-Derived Gene-Corrected Isogenic iPS Clones for Parkinson’s Disease Research. (2020). doi:10.20944/preprints202005.0001.v1

37. Krüger, R. et al. Familial parkinsonism with synuclein pathology: clinical and PET studies of A30P mutation carriers. Neurology 56, 1355–62 (2001).

38. Barbuti, P. A. et al. Generation of two iPS cell lines (HIHDNDi001-A and HIHDNDi001-B) from a Parkinson’s disease patient carrying the heterozygous p.A30P mutation in SNCA. Stem Cell Res. 48C, 101951 (2020).

39. Reinhardt, P. et al. Derivation and Expansion Using Only Small Molecules of Human Neural Progenitors for Neurodegenerative Disease Modeling. PLoS One 8, e59252 (2013).

40. Antony, P. M. A. et al. Fibroblast mitochondria in idiopathic Parkinson’s disease display morphological changes and enhanced resistance to depolarization. Sci. Rep. 10, 1569 (2020).

41. Hanss, Z. et al. Mitochondrial and Clearance Impairment in p.D620N VPS35 Patient-Derived Neurons. Mov. Disord. n/a, (2020).

42. Alam, M. & Schmidt, W. J. Rotenone destroys dopaminergic neurons and induces parkinsonian symptoms in rats. Behav. Brain Res. 136, 317–324 (2002).

43. Testa, C. M., Sherer, T. B. & Greenamyre, J. T. Rotenone induces oxidative stress and dopaminergic neuron damage in organotypic substantia nigra cultures. Brain Res. Mol. Brain Res. 134, 109–118 (2005).

44. Sulzer, D. & Surmeier, D. J. Neuronal vulnerability, pathogenesis, and Parkinson’s disease. Mov. Disord. 28, 41–50 (2013).

45. Michel, P. P., Hirsch, E. C. & Hunot, S. Understanding Dopaminergic Cell Death Pathways in Parkinson Disease. Neuron 90, 675–691 (2016).

46. Villar-Piqué, A., Lopes da Fonseca, T. & Outeiro, T. F. Structure, function and toxicity of alpha-synuclein: the Bermuda triangle in synucleinopathies. J. Neurochem. 139, 240–255 (2016).

47. Lautenschläger, J., Kaminski, C. F. & Kaminski Schierle, G. S. A-Synuclein – Regulator of Exocytosis, Endocytosis, or Both? Trends Cell Biol. 27, 468–479 (2017).

48. Vargas, K. J. et al. Synucleins regulate the kinetics of synaptic vesicle endocytosis. J. Neurosci. 34, 9364–9376 (2014).

49. Picconi, B., Piccoli, G. & Calabresi, P. Synaptic dysfunction in Parkinson’s disease. Adv. Exp. Med. Biol. 970, 553–572 (2012).

50. Schirinzi, T. et al. Early synaptic dysfunction in Parkinson’s disease: Insights from animal models. Mov. Disord. 31, 802–813 (2016).

51. Czaniecki, C. et al. Axonal pathology in hPSC-based models of Parkinson’s disease results from loss of Nrf2 transcriptional activity at the Map1b gene locus. Proc. Natl. Acad. Sci. 116, 14280 LP – 14289 (2019).

52. Burbulla, L. F. et al. Dopamine oxidation mediates mitochondrial and lysosomal dysfunction in Parkinson’s disease. Science (80-.). 357, 1255–1261 (2017).

53. Ryan, S. D. et al. Isogenic Human iPSC Parkinson’s Model Shows Nitrosative Stress-Induced Dysfunction in MEF2-PGC1α Transcription. Cell 155, 1351–1364 (2013).

54. Heman-Ackah, S. M. et al. Alpha-synuclein induces the unfolded protein response in Parkinson’s disease SNCA triplication iPSC-derived neurons. Hum. Mol. Genet. 26, 4441–4450 (2017).

55. Ryan, T. et al. Cardiolipin exposure on the outer mitochondrial membrane modulates α-synuclein. Nat. Commun. 9, 817 (2018).

56. Zanin, M. et al. Mitochondria-mitochondria interaction networks show altered topological patterns in Parkinson’s disease. bioRxiv 2020.03.09.984195 (2020). doi:10.1101/2020.03.09.984195

57. Kriks, S. et al. Dopamine neurons derived from human ES cells efficiently engraft in animal models of Parkinson’s disease. Nature 480, 547–551 (2011).

58. Kirkeby, A. et al. Generation of Regionally Specified Neural Progenitors and Functional Neurons from Human Embryonic Stem Cells under Defined Conditions. Cell Rep. 1, 703–714 (2012).

59. Stathakos, P. et al. A monolayer hiPSC culture system for autophagy/mitophagy studies in human dopaminergic neurons. Autophagy (2020). doi:10.1080/15548627.2020.1739441

60. Mertens, J., Reid, D., Lau, S., Kim, Y. & Gage, F. H. Aging in a Dish: iPSC-Derived and Directly Induced Neurons for Studying Brain Aging and Age-Related Neurodegenerative Diseases. Annu. Rev. Genet. 52, 271–293 (2018).

61. Seidel, K. et al. First appraisal of brain pathology owing to A30P mutant alpha-synuclein. Ann. Neurol. 67, 684–689 (2010).

62. van Steenoven, I. et al. Conversion between mini-mental state examination, montreal cognitive assessment, and dementia rating scale-2 scores in Parkinson’s disease. Mov. Disord. 29, 1809–1815 (2014).

