## Supplementary Figure 1 for "Gene-corrected Parkinson’s disease neurons show the A30P alpha-synuclein point mutation leads to reduced neuronal branching and function"

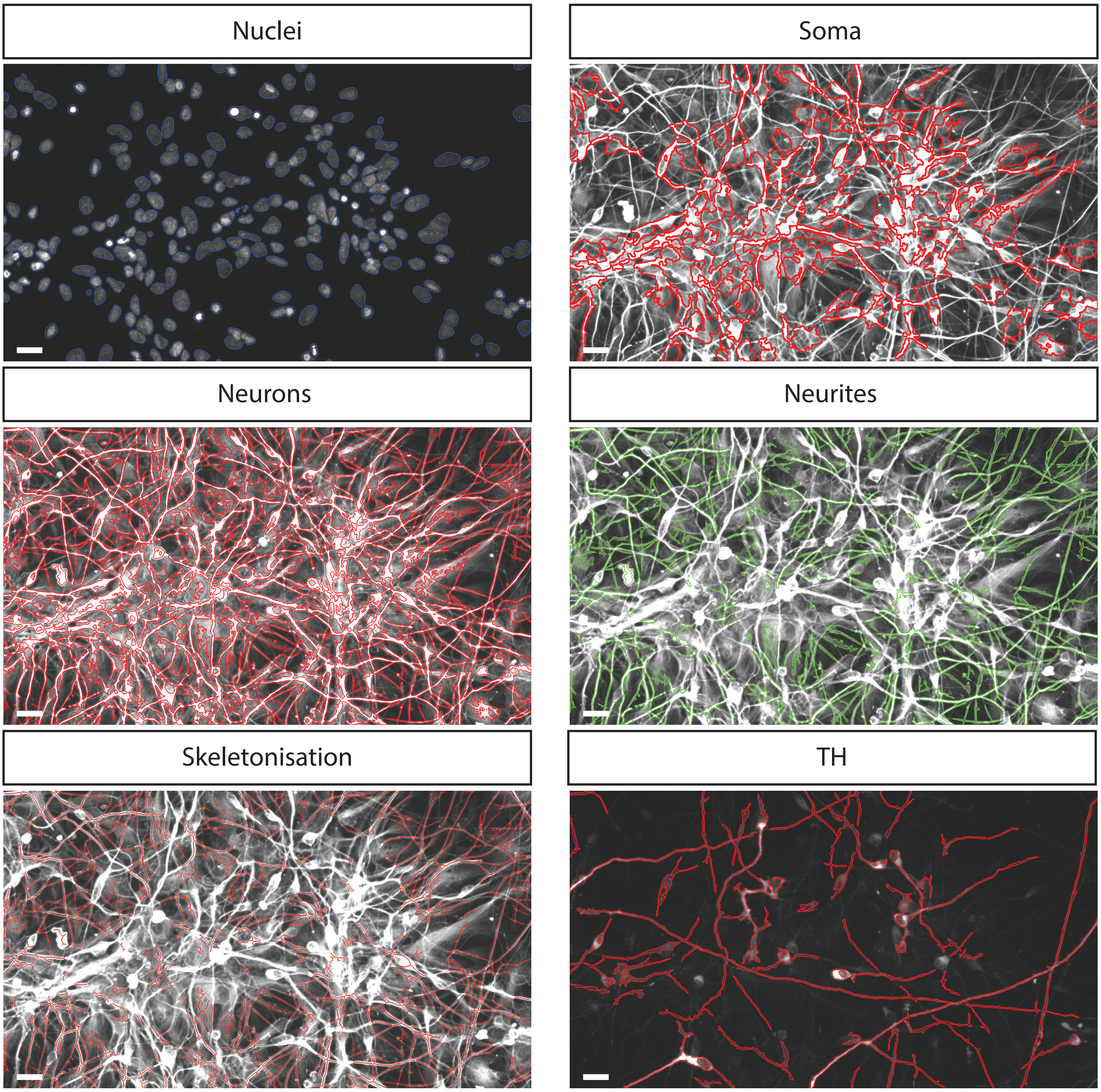


Supplementary Figure 1: Segmentation of neuronal network analysis. Representative image of a maximum intensity projection taken from a z-stack of d30 neurons fluorescently labelled with the neuronal marker Tuj1 and the dopaminergic marker TH.
