## Supplementary Figure 2 for "Gene-corrected Parkinson’s disease neurons show the A30P alpha-synuclein point mutation leads to reduced neuronal branching and function"

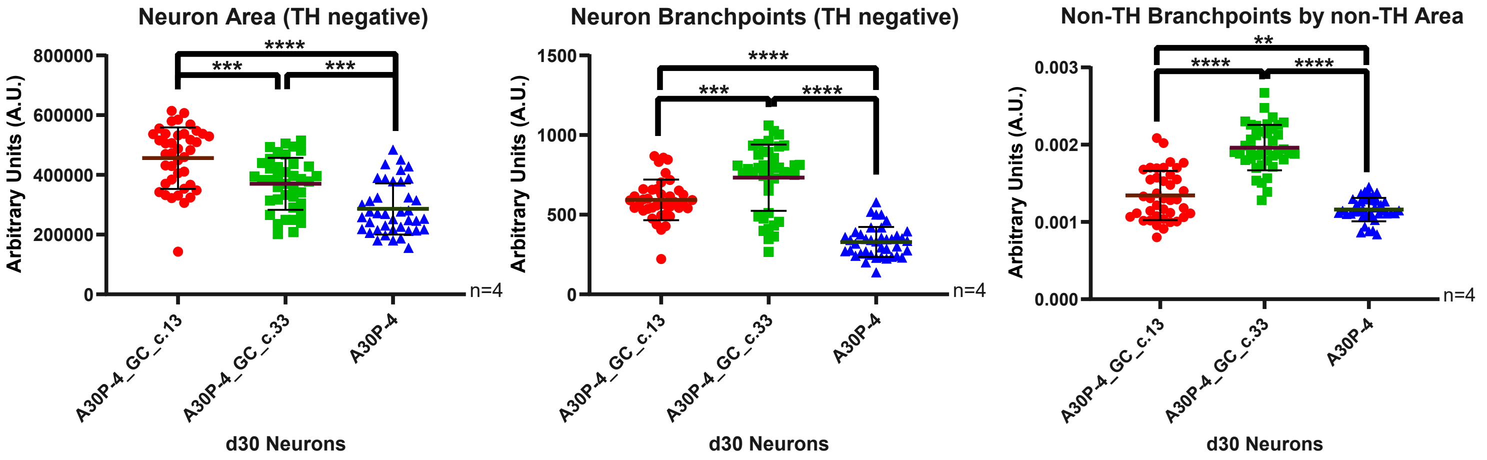


Supplementary Figure 2: Non-dopaminergic neuronal network analysis. Four biological replicates were used for the analysis with a minimum of ten Z-stacks analysed by replicate. The graphs displayed in columns show individual values that refer to the average data per Z-stack. For all statistical analyses, an ordinary one-way ANOVA was performed with Tukey's post-hoc multiple comparison test. All graphs were plotted as mean ± SD. ** $p< 0.01$, ***$p<0.001$, **** $p< 0.0001$.
